# Japanese Encephalitis Virus: A pan-proteome analysis for aggregation propensities and in vitro validation with Capsid anchor and 2K peptide

**DOI:** 10.1101/2023.09.06.556571

**Authors:** Kumar Udit Saumya, Tanya Bharadwaj, Duni C. Thakur, Deepanshu Verma, Rajanish Giri

**Author notes:** Correspondence: Dr. Rajanish Giri, School of Biosciences and Bioengineering, Indian Institute of Technology Mandi, Himachal Pradesh, 175075, India.

## Abstract

Japanese encephalitis infection is a vector-borne disease caused by the flavivirus Japanese encephalitis virus (JEV). It is responsible of severe brain infection in humans worldwide. Given the ubiquitous nature of complications and tropism associated with Japanese encephalitis (JE) infection, a holistic understanding of its molecular mechanism is essential. The phenomenon of abnormal protein aggregation into pathogenic amyloids is now increasingly linked to multiple human diseases, also known as Amyloidosis. Most are neurodegenerative disorders but amyloidosis is not restricted to a specific organ or tissue type. The overlap of viral protein aggregation with human pathologies remains limited and it is gaining momentum, especially after the devastating Covid-19 pandemic. Therefore, in this study we have examined the likelihood of aggregation for the entire collection of proteins in JEV. Multiple independent web server tools were employed to scan for potential amyloid prone-regions (APRs), and it was followed by in vitro validation using two JEV transmembrane domains, Capsid anchor and 2K peptides. These synthetic viral peptides were introduced to artificial aggregation-inducing conditions and then analyzed using a different dye-based assays and microscopy methods confirming amyloid-like fibril structure formation. We found these aggregates cytotoxic to human neuronal cell line and membrane damaging to human blood derived RBC’s. The aggregation kinetics of both peptides is enhanced in the presence of artificial membrane models and seeds of self and diabetes hallmark protein Amylin. Our findings thereby strongly suggest the possibility of JEV protein aggregation playing a vital role in its pathogenesis, opening up a broad scope of future study. Also, the interplay between JEV protein aggregation and initiation/progression of other proteopathies is possible and needs further exploration.

Graphical Abstract

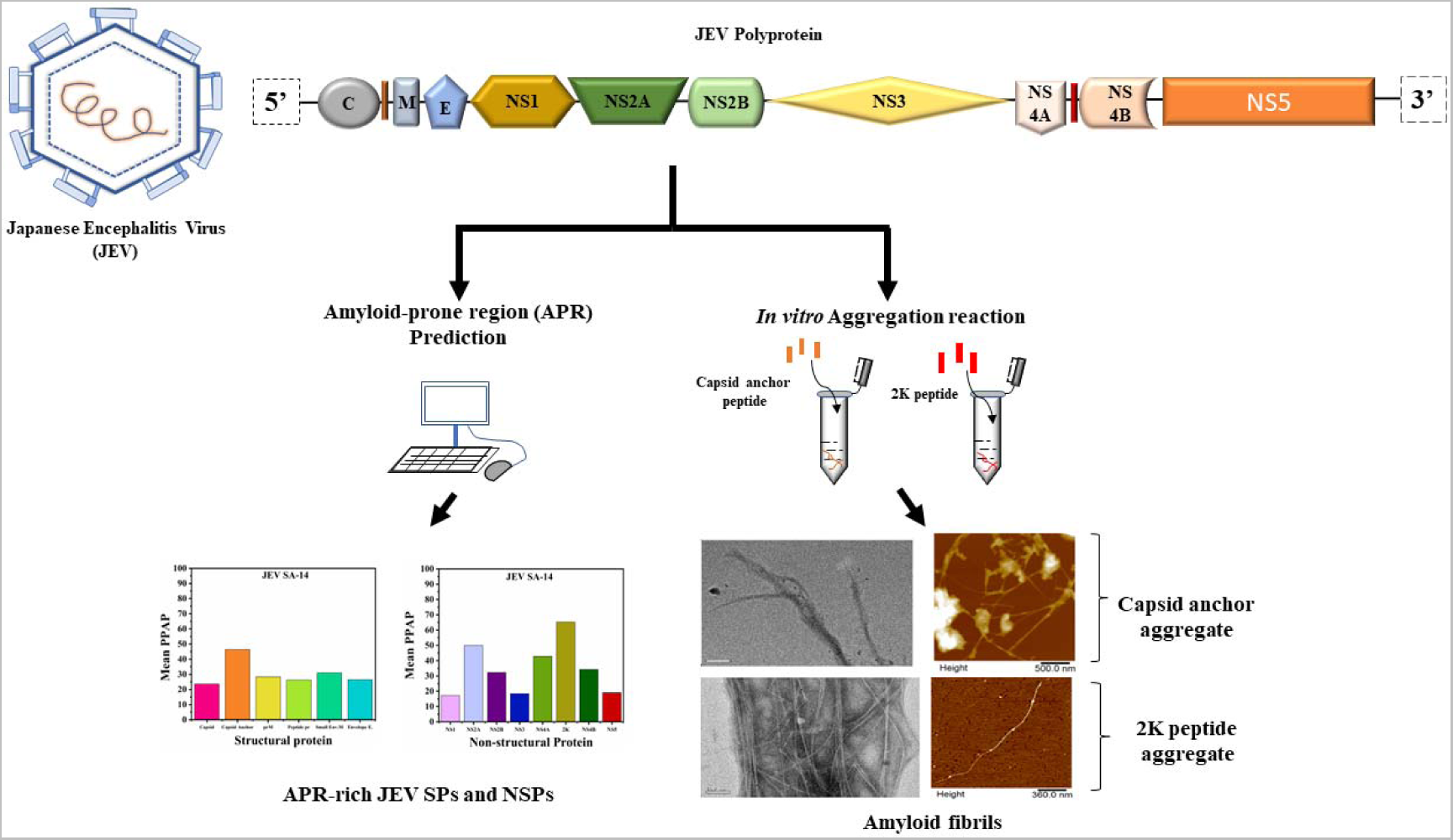

## Introduction

Japanese encephalitis virus (JEV) is an emerging arbovirus primarily spread by Culex and Aedes mosquitoes. The virus circulates between birds (natural JEV host), pigs (JEV amplifiers), and humans (JEV dead-end host) (1). It is a member of the flavivirus genus and related to the Zika virus (ZIKV), dengue virus (DENV), tick-borne encephalitis virus (TBEV), and West Nile virus (WNV) (2). In 1871 the first JEV infection was documented in Japan; since then, its geographical distribution has expanded across Asia and Australia (1). In India, JEV infection is a leading health concern, and epidemics have been reported in several states since 1955 (3). Japanese encephalitis (JE) infection causes acute viral encephalitis and various neurological illnesses in humans (4). JEV virions are icosahedral structures measuring approximately 510 Å in diameter, slightly larger than DENV2 and ZIKV. Its genome is a positive sense single-stranded RNA (Ill11kb) that encodes three structural proteins [the capsid (C), membrane (prM/M), envelope (E)], seven non-structural (NS) proteins (NS1, NS2A, NS2B, NS3, NS4A, NS4B, NS5), and two peptides namely capsid anchor and 2K (Figure 1) (5). The structural proteins (SPs) act as building blocks for new virus particle formation by primarily participating in packaging and assembly. The NSPs, on the other hand, are involved in forming the replication complex essential for viral RNA replication and innate immune response evasion (6, 7). Currently, no antiviral drugs are available against JEV. Only few vaccines have been developed to prevent infection, primarily using the live attenuated JE strain SA14-14-2, developed initially from wild-type JE SA14 (8). Therefore, combined global efforts have been ongoing for decades to enhance our understanding of the molecular mechanism involved in JEV infection and the identification of novel therapeutic targets.

**Figure 1.**
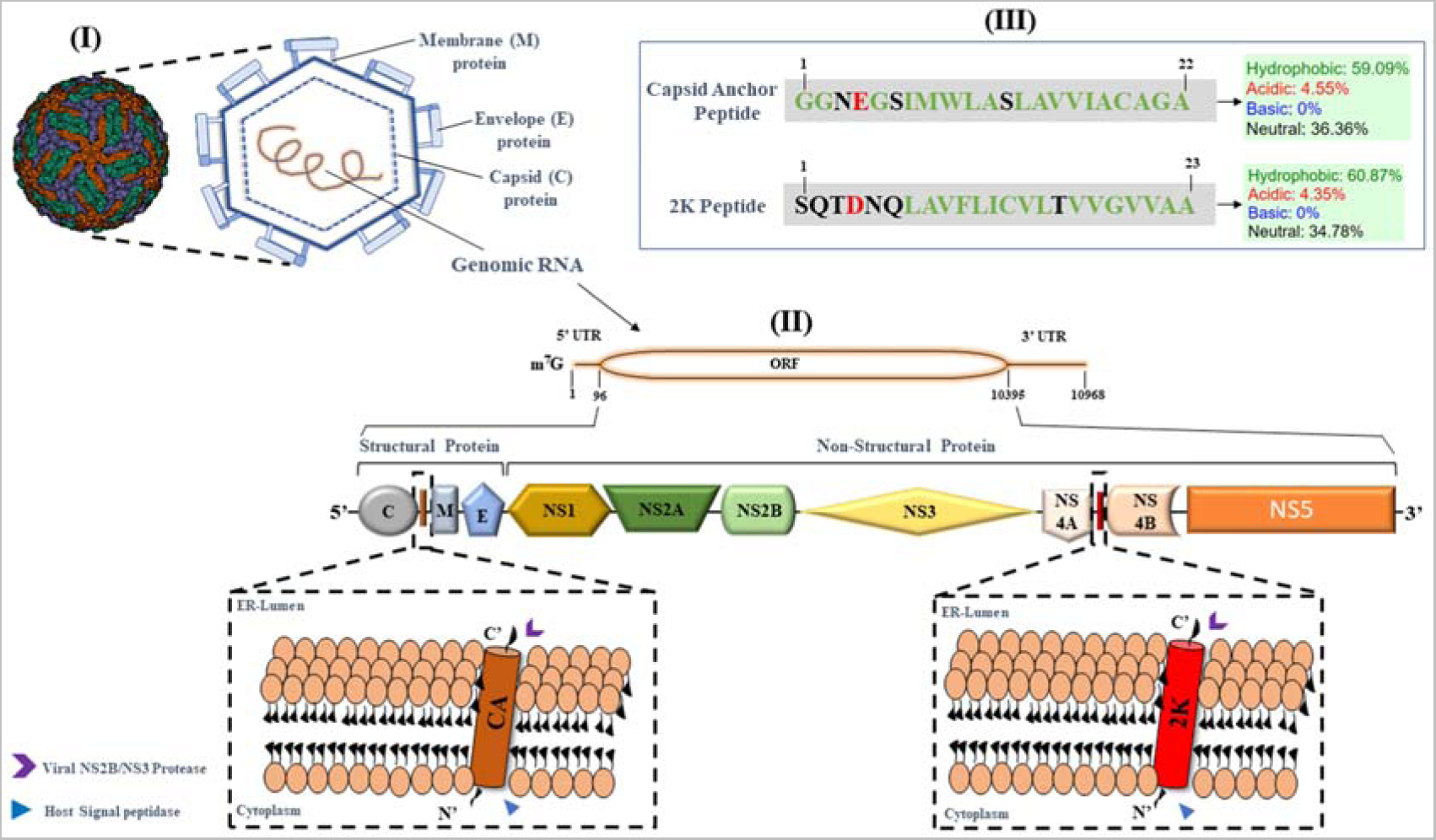
The Japanese Encephalitis (JEV) virion’s structure, genome arrangement, polyprotein composition, and sequence analysis of the Capsid anchor (CA) and 2K peptide. **(I)** Overview of JEV cryo-EM structure reported by Wang *et al.* (2017) and its schematic representation. The outer layer of the virus is formed by envelope (E) and membrane (M) proteins. The JEV genomic RNA (single-stranded, positive-sense) remains enclosed within the capsid (C) protein. **(II)** The genomic RNA is ~11kb nucleotides long. The 5’ end is a methylated cap with 95 nucleotides untranslated region (UTR). It is followed by an open reading frame (ORF) of 10,299 nucleotides and a 3’ untranslated region of 574 nucleotide residue. The open reading frame (ORF) encodes a trio of structural proteins [capsid(C), membrane (M), envelope (E)] and an array of seven non-structural proteins (NS) [NS1, NS2A,NS2B, NS3, NS4A, NS4B, and NS5]. Within the host cell cytoplasmic side, the capsid (C) protein and NS4A protein remain anchored into ER membrane by capsid anchor (CA) and 2K peptide, respectively. Topology of ER embedded CA and 2K peptide is highlighted in zoomed square boxes. During polyprotein maturation, host signal peptidase sequentially cleaves at C-2K junction and NS4A-2K junctions on the cytoplasmic side. Similarly, the viral NS2B/NS3 protease cleaves at CA-premembrane (prM) and 2K-NS4B junctions in the ER lumen side. **(III)** The sequence analysis of JEV’s Capsid anchor (CA) and 2K peptide, indicating their respective percent hydrophobicity, is represented in grey and green boxes, respectively.

The formation and accumulation of amyloid-forming proteins within and around the cells are responsible for multiple neurodegenerative and systemic diseases (9). The pathogenic proteins typically undergo misfolding into stable and highly ordered insoluble fibrillar complexes specific to the origin and associated disorder. This phenomenon is called Amyloidosis (9). Some of the well-studied diseases and their respective amyloidogenic proteins are Alzheimer’s (amyloid-beta peptides), Parkinson’s (alpha-synuclein), Huntington’s (huntingtin protein), Amyotrophic Lateral Sclerosis (TAR DNA-binding protein 43), Creutzfeldt-Jakob disease (prion protein) and type-2 diabetes (IAPP or amylin) (10–16). Interestingly, such aggregation of viral proteins and their implication in pathogenesis are also increasingly reported. A few notable examples include aggregation of glycoprotein M (EBV-gM) in Epstein-Barr virus (17), PB1 protein and nuclear exporter protein (NEP) in influenza A virus (18, 19), capsid anchor (CA) in ZIKV (20), cM45 protein in murine cytomegalovirus (21) and capsid protein in cowpea chlorotic mottle virus (CCMV) (22). In a recent investigation, it was demonstrated that proteins, signal peptides and fusion peptides from both the SARS-CoV-1 and SARS-CoV-2 viruses have the capability to form amyloid aggregates (23). Viruses exploit this phenomenon to its advances in infected host cells either directly through cytotoxicity and aberration of essential signaling pathways or indirectly by inducing misfolding of host proteins. However, the identification of APRs in viral proteins is at a very initial phase, and in-depth knowledge of the exact mechanism involved is currently lacking, but it holds exciting prospects.

In this study, therefore, we delved into assessing the aggregation propensities of the JEV proteome. Our initial computational analysis involved the identification of APR regions using multiple webserver tools. We noticed two important transmembrane domains (TMD) Capsid anchor (CA) and 2K peptide, with the highest prediction score (Figures 2 & 3). Since they are small peptides which are easy to synthesize, therefore, we continued our in vitro investigation with artificially synthesized CA and 2K peptides. These peptides were introduced to aggregation-inducing conditions, and at specific time intervals, amyloid-specific dye-based assays were used for tracking aggregation kinetics. Visualization of aggregated peptide morphology was facilitated using techniques like atomic force microscopy (AFM) and high-resolution transmission electron microscopy (HR-TEM). Further, their cytotoxic and membrane-damaging properties were assessed. Since many studies have reported that biomembrane influence aggregation behavior, morphology, and amyloid properties(24), we investigated the effect of artificial membrane models (liposomes) on the fibrillogenesis rate of JEV CA and 2K peptides. Finally, we have also demonstrated the possible cross-talk between aggregation of viral protein and Amylin, a hallmark type-2 diabetic protein.

**Figure 2.**
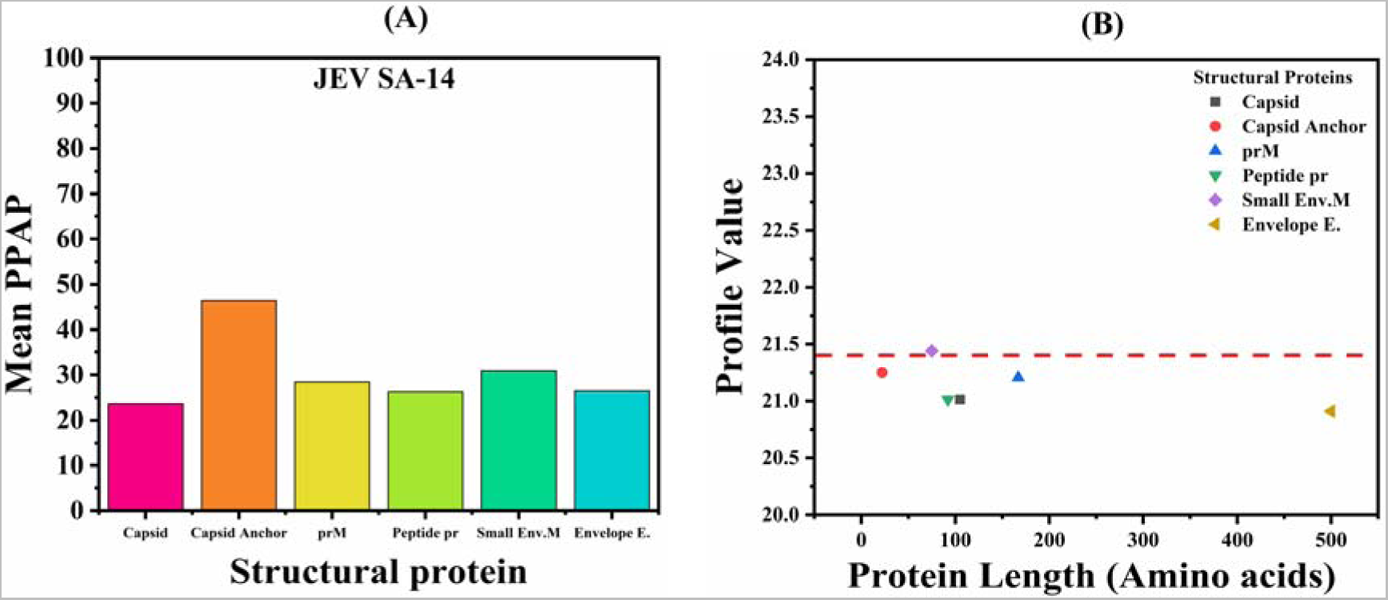
Analysis of amyloidogenic propensities in JEV Structural proteins: **(A)** Predicted percentage amyloidogenic propensity (PPAP) computed from the mean of aggregation-prone regions identified in structural proteins by prediction tools **(B)** Shows aggregation profile value of JEV structural proteins predicted by FoldAmyloid.

**Figure 3.**
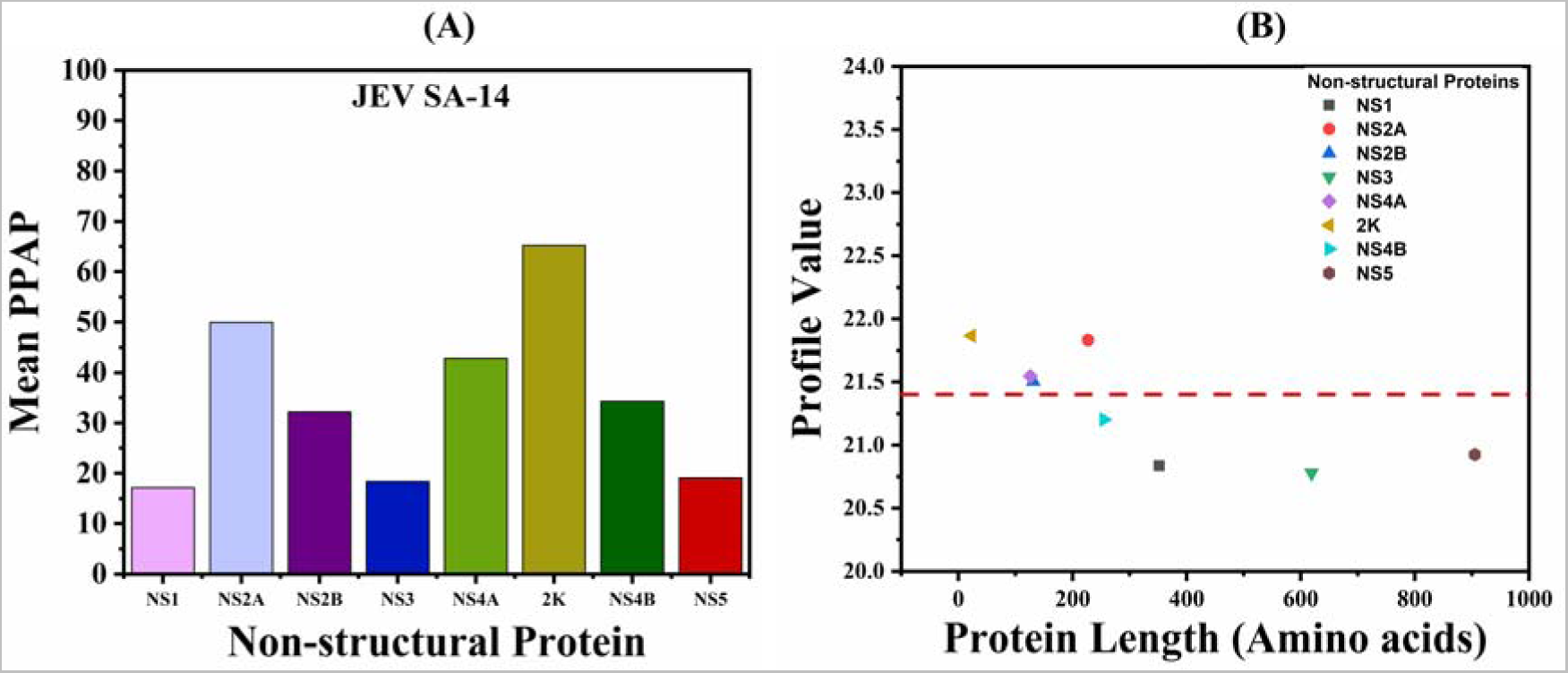
Analysis of amyloidogenic propensities in JEV Non-structural proteins. **(A)** Predicted percentage amyloidogenic propensity (PPAP) computed from the mean of aggregation-prone regions identified in structural proteins by prediction tools **(B)** Shows aggregation profile value of JEV non-structural proteins predicted by FoldAmyloid

## Methodology

### Peptide and Chemicals

The synthetic Capsid anchor peptide “GGNEGSIMWLASLAVVIACAGA” (CA; 22 amino acid residues; purity 90.5%) and 2K peptide “SQTDNQLAVFLICVLTVVGVVAA” (23 amino acid residues; 95.9% purity) were procured from Genscript, USA. The certificate of peptide purity is attached in the supplementary file (Figure S1-S3). Additional chemicals, reagents, fluorescent dyes, negative staining agent, etc. were ordered from Sigma Aldrich, St. Louis, USA. Copper girds and mica sheets were sourced from TED PELLA INC., USA. The neutral lipid 1,2-dioleoyl-sn-glycero-3-phosphocholine (DOPC) and negatively charged lipid 1,2-dioleoyl-sn-glycero-3-phospho-L serine (DOPS) were acquired from Avanti Polar Lipids (Alabaster, AL, U.S.A.). For cell culture, chemicals and reagents were procured from GibcoTM. The human neuroblastoma cell line (SH-SY5Y) was used for cytotoxicity assays.

### Prediction of amyloidogenic regions in JEV proteome

The JEV proteome sequences corresponding to the SA-14 strain (UniProt accession ID: P27395) were retrieved from the UniProt database. For amyloid-prone region (APR) identification, a total of 5 different tools were used: Metamyl (25), FoldAmyloid (26), Tango (27), Aggrescan (28), and FISH Amyloid (29). Complete default parameters were used in all tools except for Tango, in which the pH was set to 7.4 and temperature to 37°C. Briefly, The meta-prediction tool MetAmyl combines the functions of four individual tools namely SALSA, PAFIG, Foldamyloid and Waltz to select input sequence fragments with aggregation probability (25). FoldAmyloid assesses the likelihood of backbone hydrogen bond formation to classify amyloidogenic peptides. A threshold of 21.4 is assigned by the server as the cut-off score (26). The algorithm in TANGO is designed to predict hydrophobic beta-sheets using principles of secondary structure formation (27). Aggrescan relies on an aggregation propensity scale developed from in vivo experimental results for natural amino acid residues. It also operates under the assumption that concise and specific sequences play a role in regulating protein aggregation (28). FISH Amyloid functions on machine learning prediction method that has been trained on recognizing co-occurrence patterns of residues in sequences (29).

### Preparation for the aggregation assays

CA, 2K and human amylin (hIAPP) peptides upon removal from −80□ were first placed on ice for 10 minutes and then used for weighing. To eliminate any pre-existing aggregates the peptides were first dissolved in 400 µl of 100% HFIP (1,1,1,3,3,3-hexafluoro-2-propanol) was used to dissolve the peptides. A brief vacuum spin was performed to evaporate HFIP and get dry peptides. The Genscript recommended analytical grade DMSO (20%) was used as a solvent to resuspend peptides and then transferred to sodium phosphate buffer (20mM; pH 7.4) to acquire the desired final concentration. After dissolving the peptides, samples were transferred to incubation at 37°C while being continuously stirred (1000 rpm) using the Eppendorf ThermoMixer C. The stock peptide concentration used for dye-based assays (ThT, ANS, CR), imaging (TEM, AFM), cytotoxicity (MTT), and hemolysis assay was 1mg/ml & 3mg/ml. For ThT kinetics, the final peptide concentration for both peptides were kept at 0.05mg/ml uM. Seeds of desired concentration were prepared by probe sonication of aggregated samples for 10 min at 25 amplitude on ice to avoid heating of samples(30).

### HR-TEM and AFM imaging

The HR-TEM images were obtained using FP 5022/22-Tecnai G2 20 S-TWIN, FEI. A twenty-fold dilution of the aggregated peptide was done and then added to carbon-coated copper grids (200-mesh; Ted Pella, Inc, USA) through drop-casting technique. A freshly prepared 3% ammonium molybdate solution was used for negative staining of TEM grids. The grids with sample on it were allowed to dry overnight at room temperature.

Tapping-mode atomic force microscope (AFM) (Dimension Icon system from Bruker) was used to visualize aggregates. The measurements were conducted by adding a thirty-fold diluted solution of aggregated samples onto clean mica surface. Incubation of sample on mica sheet was done for an hour following which the sheet surfaces were rinsed with deionized water. Mica sheets with sample on it were allowed to dry at room temperature overnight, and images were then captured. TEM and AFM images were acquired using CA and 2K peptides that was kept under aggregation incubation for 240 h and 96 h, respectively.

### Liposome & seed preparation

A thin-filmed hydration method was used for preparing liposomes/large unilamellar vesicles (LUVs) (31). Lyophilized DOPC was weighed and dissolved in analytical grade chloroform solution, whereas pre-dissolved DOPS in chloroform was already available. These lipids were introduced into two separate round bottom flasks (RB), and the chloroform was eliminated under vacuum at room temperature. To ensure complete removal of the solvent, the RB flasks were placed in a vacuum within a desiccator overnight. Subsequently, sodium phosphate buffer (20mM, pH 7.4) was administered to re-hydrate the dry lipid films and transferred to a mini-extruder setup. In essence, a sequence of three freeze-thaw cycles involving liquid nitrogen and water bath (set at 60°C) was executed, each accompanied by intermittent vortexing for 2 mins. This was followed by extrusion through the mini extruder (Avanti Polar Lipids, Inc. USA) fitted with 100 nm polycarbonate membrane cut-off filter for a total 30 times. The final lipid concentration of the liposomes were 60.42 mM and 12.34 mM for DOPC & DOPS, respectively. They were stored in sterile amber colored ampules at 4°C and used within three days. The homogeneity in size was assessed via Dynamic Light Scattering (DLS) using a Zetasizer Nano ZS (Malvern Instruments Ltd., UK) equipped with a laser (633 nm) positioned at an angle of 173°.

### Cell viability assay

The viability of SH-SY5Y neuroblastoma cells when exposed to aggregates of CA and 2K was evaluated using a colorimetric assay. This assay employs the reduction of a yellow tetrazolium salt 3-(4,5-dimethylthiazol-2-yl)-2,5-diphenyltetrazolium bromide (MTT), by metabolically active cells to produce purple formazan crystals. Dulbecco’s modified Eagle’s medium/Nutrient Mixture F-12 Ham supplemented with fetal bovine serum (10% v/v) was used to culture the cells in 25cm^2^ T-flasks (Thermo Scientific). CA aggregates and 2K peptide aggregates were prepared by incubating monomeric peptides in aggregation inducing condition for 240 h and 96h, respectively. The aggregates were then washed in sterile PBS buffer thrice to remove DMSO and finally resuspended in the DMEM media. The SH-SY5Y cells were seeded in a 96-well plate (hemocytometer count 6000/well) and allowed to completely adhere on plate surface for 24 h. The cells were treated with varying concentrations of CA and 2K aggregate samples ranging from 5-500µM then incubated for 72 h at 37°C in CO_2_ incubator. Next, freshly prepared 10µl MTT at 0.5 mg/ml final concentration was added to each well. The 96-well plate was then incubated for 3 h at 37°C. The treatment was stopped by washing out MTT and analytical grade DMSO was added to dissolve the formazan crystals. Final absorbance reading was acquired using TECAN Infinite M200 PRO multimode microplate reader. Absorbance at wavelength 570 nm (reference wavelength at 630 nm) was used. The data were collected in triplicates for each concentration and percent viability with respect to control (PBS treated) was calculated.

### Hemolysis assays

Hemoglobin release into the plasma as a result of red blood cell (RBC) lysis was monitored following incubation with the same JEV CA and 2K aggregates mentioned above. The experimental protocol was approved by the Institution’s (IIT-Mandi) Ethical Committee (approval letter number: IIT Mandi/IEC (H)/2020/17th January/P1). Blood samples were collected in EDTA treated vials from healthy volunteers with their consents at IIT-Mandi health center. The blood samples were immediately used for RBC isolation. Briefly, 2mL blood sample was centrifuged (1,500 rpm for 10 min) and pellet was washed thrice with 20mM phosphate-buffered saline (150 mM NaCl, pH 7.4). After final wash the pellets were resuspended at 10% v/v suspension of RBCs and PBS. Exactly, 200 μL of this suspension was transferred to sterile mini centrifuge tubes (1.5mL), followed by addition of CA and 2K aggregate fibril at various concentrations (ranging 5–250 μM). The plate was placed on a shaker (set at 300 rpm; 37°C) and incubated for a duration of 3 h. After the incubation period was over, the samples were subjected to 10 mins of centrifugation (at 1600 rpm). Supernatant was carefully collected and added to a 96-well plate. Finally, absorbance (at 540nm) was measured using a multimode microplate reader (TECAN Infinite M200 PRO). Measurements in triplicates were recorded for each concentration of aggregate. The controls used in this reaction setup were PBS treated (no hemolysis) and Triton-X100 (100% hemolysis). The hemolytic activity of the CA fibrils was determined in percentage using the following Equation.

Percentage of hemolysis = (AT-AC)/(AX-AC)*100

Where,

AT = Absorbance for supernatant for aggregate treated RBCs

AC= Absorbance for supernatant of PBS treated RBSs, and

AX= Absorbance for supernatant of 1% Triton X-100 treeated RBCs.

## Results & Discussion

### 1. Prediction of aggregation-prone regions in JEV proteome

It is known that proteins typically possess a natural/inherent tendency to form pathogenic or non-pathogenic aggregates due to the presence of amyloidogenic sequences within them, also known as amyloid-prone regions (APRs)(32). These APRs’ propensity to induce aggregation depends on critical features such as sequence, charge, secondary structure, and hydrophobicity(33). In our study, five open-source predictors, namely Metamyl (25), FoldAmyloid(26), TANGO (27), Aggrescan(28), and FISH(29) were used. From our initial analysis, we found APRs spread throughout JEV proteome (Table 1 & 2), and the calculated mean predicted percentage amyloidogenic propensity (PPAP) was comparatively higher in NSPs (Figures 2 & 3).

**Table 1.** Scores/values of aggregation propensities predicted in JEV Structural proteins using five different web server tools (MetAmyl, FoldAmyloid, TANGO, Aggrescan, FISH).

| JEV PROTEIN | AMYLOID PREDICTORS |  |  |  |  |
| --- | --- | --- | --- | --- | --- |
|  | MetAmyl | FoldAmyloid | TANGO | Aggrescan | FISH |
| CAPSID (1-105) | 23-38, 43-55, 70-77, 81-86, 90-95 | 23-29, 32-37, 45-58 | 33-38,46-54 | 27-37,44-59,89-94 | - |
| CAPSID ANCHOR (106-127) | 11-20 | 7-18 | 7-20 | 6-20 | - |
| prM (128-294) | 10-15, 18-27, 34-43, 48-53, 71-79,93-101,124-129,151-160 | 10-14, 21-25, 33-39, 41-45, 66-70, 74-78, 126-130, 134-139, 141-145, 153-162 | 74-78,133-146,153-162 | 8-14,20-25,36-44,75-79,134-146,152-163 | 3-7,21-26,35-39,75-79,124-128,153-158 |
| Peptide pr (128-219) | 10-15, 18-27, 34-43, 48-53, 71-79 | 10-14, 21-25, 33-39, 41-45, 66-70, 74-78 | 74-78 | 8-14,20-25,36-44,75-79, | 3-7,21-26,35-39,75-79 |
| Small envelope protein M (220-294) | 1-9, 32-37, 59-68 | 34-38, 42-47, 49-53, 61-70 | 41-54,61-70 | 34-38,42-54,60-71 | 32-36,61-66 |
| Envelope protein E (295-794) | 12-17, 19-26, 28-36, 38-48, 56-73, 86-95, 118-126, 138-148, 155-172, 182-187, 198-215, 247-258, 266-283, 317-328, 345-345, 350-365, 367-374, 379-388, 405-412,428-448, 455-460, 479-498 | 1-5, 19-25, 42-48, 58-63,139-144,200-205,211-221,237-241,322-326,337-341,457-461,465-472,480-486,490-496 | 139-143,200-207,265-272,339-343,381-386,463-473,479-496 | 19-30,55-61,63-73,108-112,137-146,178-183,198-218,265-283,303-308,319-328,339-347,351-358,365-370,379-383,404-408,428-442,444-473,480-500 | 42-46,64-68,152-156,210-214,320-325,432-436,472-476,489-493 |

#### 1.1 Abundance of APRs in JEV structural proteins (SPs)

The amino terminus of the flavivirus genome encodes three structural proteins, among which capsid (C) is translated first, followed by two glycoproteins: membrane (M) expressed as prM, a precursor to M (34), and envelope protein (E). Each glycoprotein contains two transmembrane helices, and during particle maturation, proteolytic cleavage yield pr peptide and M protein. The capsid is involved with packaging the viral genome, nucleocapsid core formation, assembly, and lifecycle (35). JEV C comprises four alpha helices (α1-α 4) where all except α3 are involved in capsid dimerization(36). We observed APRs of different lengths encompassing all four α-helices. Immature capsid remains anchored into ER membrane through its C-terminal domain (residues 106-127) called capsid anchor (CA)(36). During the maturation of the capsid, the CA is cleaved-off, and mature capsid protein is released into the cytoplasm (36). Since the membrane-embedded CA performs multiple essential functions in flaviviruses, we used the CA sequence (22 residues) separately for our analysis. Our results show the JEV CA sequence with the highest mean aggregation propensity among all three structural proteins (Figure 2A), especially residues 7-20, commonly overlapping between three of the five prediction tools. A similar approach was followed for the remaining structural proteins. The uncleaved prM act as a chaperone for E protein folding and might play a role in immune evasion(34). Cellular attachment and virus-host membrane fusion takes is mediated by the E protein(37). Their predicted APRs are shown in Table 1. Briefly, the mean PPAP of M and E protein was marginally higher (>24%) than capsid. In contrast, small envelope protein M, responsible for budding(38), is the only SP above the threshold profile value predicted by Fold amyloid.

#### 1.2 Abundance of APRs in JEV non-structural proteins (NSPs)

We found several APRs interspersed within the protein sequence of all seven JEV non-structural proteins (NSPs) (Figure 3 & Table 2). The first NSP NS1 is involved in crucial steps of viral infection cycle. It recruits other NSPs for replication complex formation and antagonizes the host complement system as an immune evasion strategy(39, 40). Our analysis shows NS1 with the lowest mean PPAP. Predicted common potential APRs within the N-terminal and central domains are regions reportedly involved in crucial interactions with viral and host-binding partners(39). NS1 is succeeded by NS2A and NS2B proteins. Flavivirus NS2A is a hydrophobic protein and its primary role may involve synthesis of viral RNA, expression regulation of upstream NS1 protein and inhibition of host type-I interferon response(41). NS2B on the other hand is involved in viroporins formation, membrane destabilization and acts as cofactor for downstream NS3 protease(42, 43). Both have multiple APRs according to our prediction results and constitute two of the four NSPs above the FoldAmyloid threshold profile value (Figure 3B).

**Table 2.**
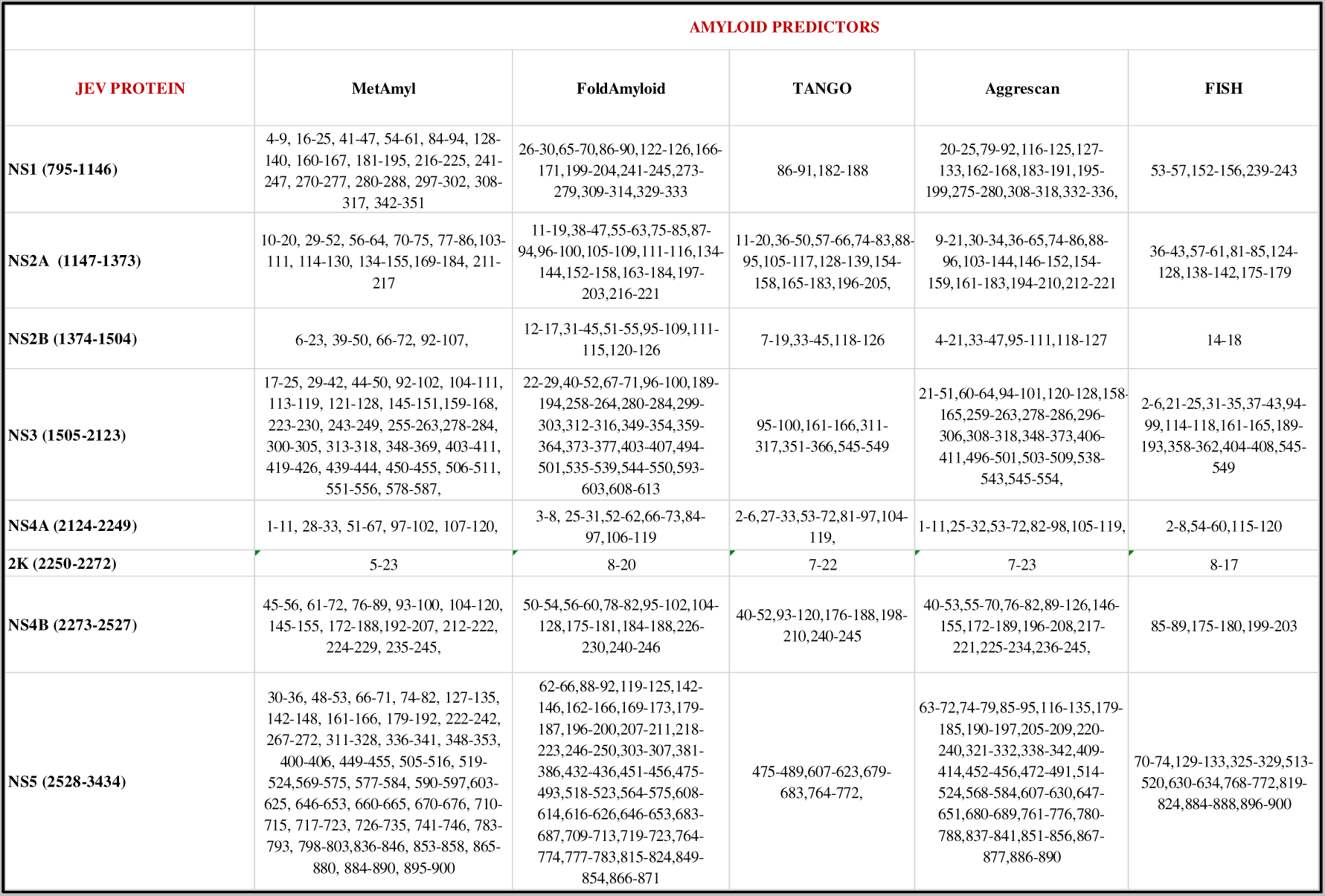
Scores/values of aggregation propensities predicted in JEV Structural proteins using five different web server tools (MetAmyl, FoldAmyloid, TANGO, Aggrescan, FISH)

| JEV PROTEIN | AMYLOID PREDICTORS |  |  |  |  |
| --- | --- | --- | --- | --- | --- |
|  | MetAmyl | FoldAmyloid | TANGO | Aggrescan | FISH |
| NS1 (795-1146) | 4-9, 16-25, 41-47, 54-61, 84-94, 128-140, 160-167, 181-195, 216-225, 241-247, 270-277, 280-288, 297-302, 308-317, 342-351 | 26-30,65-70,86-90,122-126,166-171,199-204,241-245,273-279,309-314,329-333 | 86-91,182-188 | 20-25,79-92,116-125,127-133,162-168,183-191,195-199,275-280,308-318,332-336, | 53-57,152-156,239-243 |
| NS2A (1147-1373) | 10-20, 29-52, 56-64, 70-75, 77-86,103-111, 114-130, 134-155,169-184, 211-217 | 11-19,38-47,55-63,75-85,87-94,96-100,105-109,111-116,134-144,152-158,163-184,197-203,216-221 | 11-20,36-50,57-66,74-83,88-95,105-117,128-139,154-158,165-183,196-205, | 9-21,30-34,36-65,74-86,88-96,103-144,146-152,154-159,161-183,194-210,212-221 | 36-43,57-61,81-85,124-128,138-142,175-179 |
| NS2B (1374-1504) | 6-23, 39-50, 66-72, 92-107, | 12-17,31-45,51-55,95-109,111-115,120-126 | 7-19,33-45,118-126 | 4-21,33-47,95-111,118-127 | 14-18 |
| NS3 (1505-2123) | 17-25, 29-42, 44-50, 92-102, 104-111, 113-119, 121-128, 145-151,159-168, 223-230, 243-249, 255-263,278-284, 300-305, 313-318, 348-369, 403-411, 419-426, 439-444, 450-455, 506-511, 551-556, 578-587, | 22-29,40-52,67-71,96-100,189-194,258-264,280-284,299-303,312-316,349-354,359-364,373-377,403-407,494-501,535-539,544-550,593-603,608-613 | 95-100,161-166,311-317,351-366,545-549 | 21-51,60-64,94-101,120-128,158-165,259-263,278-286,296-306,308-318,348-373,406-411,496-501,503-509,538-543,545-554, | 2-6,21-25,31-35,37-43,94-99,114-118,161-165,189-193,358-362,404-408,545-549 |
| NS4A (2124-2249) | 1-11, 28-33, 51-67, 97-102, 107-120, | 3-8, 25-31,52-62,66-73,84-97,106-119 | 2-6,27-33,53-72,81-97,104-119, | 1-11,25-32,53-72,82-98,105-119, | 2-8,54-60,115-120 |
| 2K (2250-2272) | 5-23 | 8-20 | 7-22 | 7-23 | 8-17 |
| NS4B (2273-2527) | 45-56, 61-72, 76-89, 93-100, 104-120, 145-155, 172-188,192-207, 212-222, 224-229, 235-245, | 50-54,56-60,78-82,95-102,104-128,175-181,184-188,226-230,240-246 | 40-52,93-120,176-188,198-210,240-245 | 40-53,55-70,76-82,89-126,146-155,172-189,196-208,217-221,225-234,236-245, | 85-89,175-180,199-203 |
| NS5 (2528-3434) | 30-36, 48-53, 66-71, 74-82, 127-135, 142-148, 161-166, 179-192, 222-242, 267-272, 311-328, 336-341, 348-353, 400-406, 449-455, 505-516, 519-524,569-575, 577-584, 590-597,603-625, 646-653, 660-665, 670-676, 710-715, 717-723, 726-735, 741-746, 783-793, 798-803,836-846, 853-858, 865-880, 884-890, 895-900 | 62-66,88-92,119-125,142-146,162-166,169-173,179-187,196-200,207-211,218-223,246-250,303-307,381-386,432-436,451-456,475-493,518-523,564-575,608-614,616-626,646-653,683-687,709-713,719-723,764-774,777-783,815-824,849-854,866-871 | 475-489,607-623,679-683,764-772, | 63-72,74-79,85-95,116-135,179-185,190-197,205-209,220-240,321-332,338-342,409-414,452-456,472-491,514-524,568-584,607-630,647-651,680-689,761-776,780-788,837-841,851-856,867-877,886-890 | 70-74,129-133,325-329,513-520,630-634,768-772,819-824,884-888,896-900 |

The flaviviral NS3 protein is a tri-functional protein with serine protease, NTPase, and helicase activity(42, 44). The N-terminal is the protease domain, while its C-terminal is involved in helicase activity. Results from Metamyl, Aggrescan, FISH, and FoldAmyloid predicted common APRs in both of these functional domains. Very little information is available about the small hydrophobic protein NS4A that remains connected to the larger NS4B protein through the 2K peptide (NS4A-2K-NS4B). Structurally, the 126 residues long NS4A protein consists of a hydrophilic N-terminal domain and three ER lumen-embedded hydrophobic regions (pTMS1–pTMS3)(45). Flaviviral NS4A essentially regulates the NS3 helicase activity(46). Our analysis found almost common APRs in the N-terminal portion, facilitating NS4A homo-oligomerization. Mutagenesis analysis of two NS4A residues, Y3N and A97E(47), was previously reported to revert replication defects in mutated JEV (NS4A-K97R). These two residues were also commonly found within APRs by all five-prediction tools. A short-transmembrane peptide 2K connects NS4A and NS4B in the polyprotein. According to our prediction, the 2K peptide has the highest mean PPAP (>65%) among all SPs and NSPs. It functions as a signal sequence for NS4B protein translocation into ER lumen(48). The 2K peptide is dissociated from NS4A and NS4B proteins through flaviviral protease cleavage at the NS4A-2K site, followed by host signalase cleavage at the 2K-NS4B site(48). The NS4B consists of 2 transmembrane domains at the N-terminal and three more at its C-terminal region. Our prediction analysis found APRs mostly in their C-terminal transmembrane portions. Lastly, NS5 is the most conserved and the largest in the flaviviral polyprotein. Residues 1-265 constitutes the methyltransferase (Mtase) domain involved in the 5’methyl capping of RNA, while the 276-905 residues making up central and C-terminal regions functions as RNA-dependent RNA polymerase enzyme(49). Besides, NS5 also blocks cellular antiviral response via inhibition of interferon-alpha/beta signaling (50). The predicted APRs in both these vital enzymatic domains of NS5 are listed in Table 2, except for TANGO (APR predicted only in the RDRP domain).

### 2. Experimental validation of predicted aggregation-prone JEV Capsid anchor and 2K peptides

Subsequent to our in-silico examination of the JEV protein sequences, we proceeded to explore the aggregation tendencies of synthetically produced CA and 2K peptides. To achieve this, we employed a blend of dye-based assays (ThT, ANS, CR) and high-resolution microscopy techniques (HR-TEM, AFM) in order to monitor the rate of fibril formation and observe the structural characteristics of the formed fibrils. In addition, artificial membrane models/liposomes carrying different charges were also prepared to assess whether they influenced the fibrillation rate. Finally, cytotoxic and membrane-damaging properties of CA and 2K peptide aggregates were assessed through MTT and RBC hemolysis assays.

#### 2.1 Dye-based assays and morphological characterization of JEV Capsid anchor

The JEV CA is a short membrane-embedded peptide of 22 amino acid residues(36). In flavivirus, the canonical role of CA is to anchor immature capsid into ER membrane while its hydrophobic C-terminus act as a signal sequence for the translocation of prM protein into ER lumen(51, 52). We have shown above that this CA C-terminal region is predicted as highly amyloidogenic. Moreover, CA assumes a crucial role in upholding the stability of envelope proteins, the assembly process of viral proteins, and maturation of new viral particles(53). Our previous study also reported ZIKV CA peptide forms cytotoxic amyloid fibrils(20). Based on this information and our preliminary findings, we were intrigued to study CA peptides at physiological aggregation conditions.

We used three amyloid-specific dyes to monitor the aggregation behavior of the JEV CA peptide. These dyes [ThT (thioflavin T), ANS (4,4’-dianilino-1,1’-binaphthyl-5,5′-disulfonic acid, dipotassium salt), CR (Congo red)] have been previously reported for studying in vitro aggregation of viral proteins(20, 23). The ThT dye shows a strong fluorescence signal upon binding with beta sheet-rich structures, which are abundant in amyloid fibrils(54). Our findings revealed approximately ~6-fold increase in ThT fluorescence upon aggregation (Figure 4A). ANS is a fluorescent dye routinely used to detect intermediates during protein conformational changes(55). This dye has a high affinity for hydrophobic binding pockets on aggregating protein surfaces. We found that the JEV CA aggregate was ANS positive, and the observed characteristic red shift of 42nm (from 530 nm to 488 nm) indicates probable aggregation-induced conformational change (Figure 4B). The anionic CR dye, on the other hand, binds to beta sheet-rich amyloid proteins and exhibits prominent dichroism under polarized light(56). Our CR assay with CA peptide exhibited a slight red shift in absorbance spectra from 491 nm to 495 nm (Figure 4C). These results suggest JEV CA peptide aggregates into β sheet-rich structures in physiological pH and temperature.

**Figure 4.**
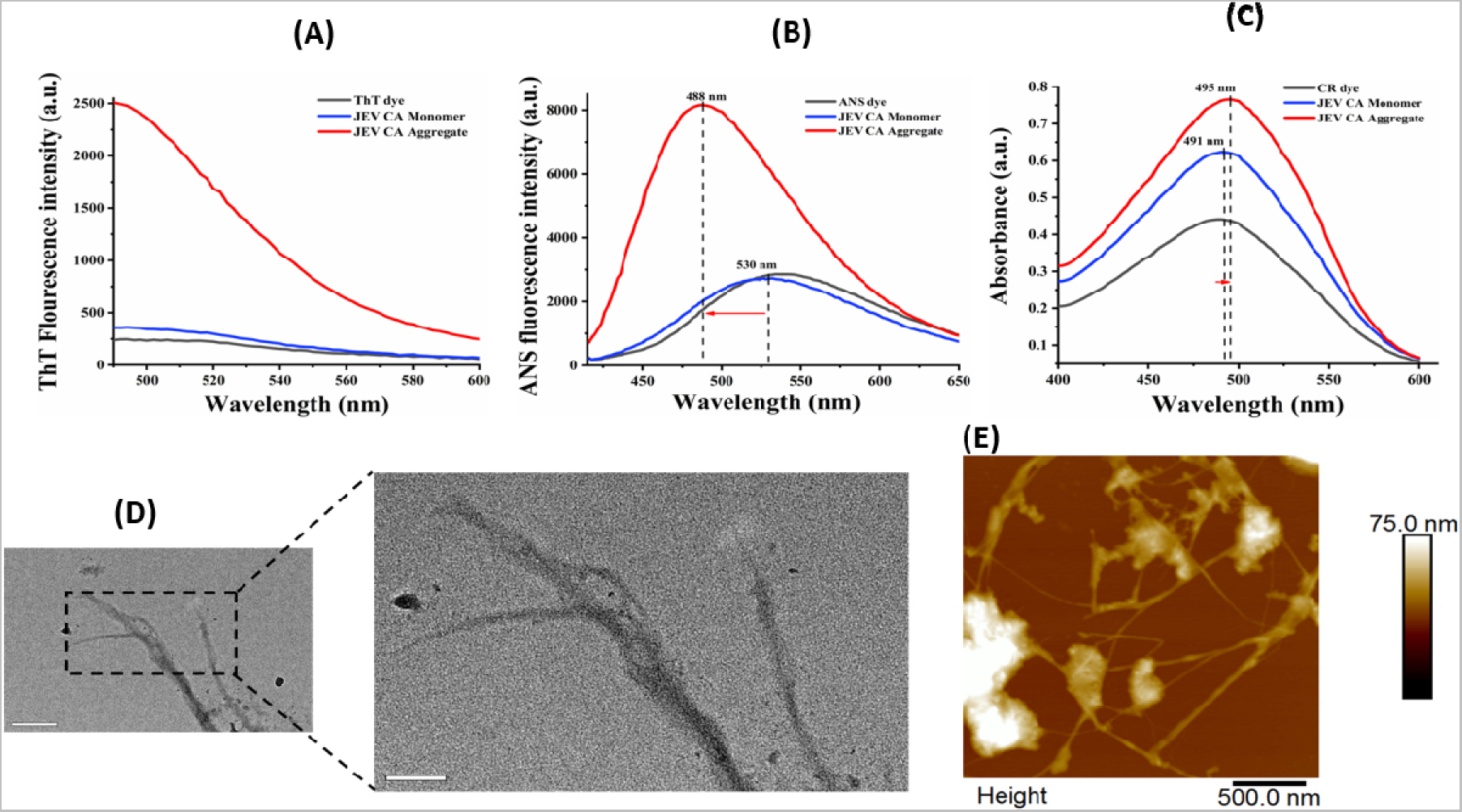
In vitro aggregation of the JEV CA peptide. **(A)** Scans of ThT fluorescence spectra performed with CA peptides at 1mg/ml concentration showing increased fluorescence intensity **(B)** ANS fluorescence emission spectra scan showing blue-shift **(C)** Absorption spectra scan of congo red dye bound capsid anchor peptides samples measured showing red-shift. TEM and AFM images of CA aggregates display amyloid fibril-like structures with dense protruding.

Next, HR-TEM and AFM imaging were conducted attain a deeper understanding of the morphological attributes present in the CA aggregated samples. We observed amyloid-like aggregates with densely interconnected fibrils of varying lengths. Similar morphology has been reported for ZIKV CA aggregates and several other viral and non-viral amyloid proteins(20, 23, 57).

#### 2.2 Dye-based assays and morphological characterization of JEV 2K peptide

In JEV, its non-structural proteins NS4A and NS4B remain connected through a short 23 amino acid transmembrane peptide called 2K(58). During maturation, the NS4A-2K-NS4B complex is sequentially cleaved off by host signal peptidase and viral NS2B-NS3 protease, releasing mature NS4A and NS4B proteins. The 2K peptide is crucial for the stable maturation of NS4B as it regulates its correct folding and insertion into ER membrane(59, 60). Flavivirus infection is known to induce the restructuring of cytoplasmic membranes for efficient viral replication. In WNV, deleting its 2K domain leads to less prominent membrane rearrangement(61). However, no information is available on 2K peptide aggregation in isolation.

Using different computational tools, we have shown that JEV 2K peptide shows a high aggregation tendency. We observed in 2K aggregate 7-fold increase in ThT fluorescence against its monomeric form. An increase in ANS fluorescence intensity accompanied by a blue-shift of 5nm indicates a conformational transition of 2K monomers towards an aggregation state. Our CR assay revealed a 4nm red-shift in absorbance spectra of 2K, similar to CA results. Thus, JEV 2K peptide was positive for all three amyloid-specific dyes. Next, TEM and AFM images of 2K aggregates revealed long networked amyloid-like fibrillar structures. We quantitively measured the width of the fibril captured through TEM at three different locations and found them to be 4.24nm, 4.42nm and 4.57nm.

#### 2.3 Fibrillogenesis is enhanced by membrane-mimicking Liposomes and seeds derived from aggregates of viral peptide and Amylin protein

Environmental conditions strongly influence protein aggregation. Any temperature, pH, or ionic strength changes lead to polymorphic amyloid fibrillation(24). Biomembrane are factors that regulate amyloidogenesis by lowering the lag time and inducing protein conformational changes(24, 62). Fundamental properties of a biomembrane, such as its components, lipid type, size, and curvature, regulate the aggregation behavior, structure, and even biochemical properties(24). Since CA and 2K peptides are transmembrane domains (TMDs) that remain embedded in the host cell ER membrane, we investigated the effect of membrane-mimicking models on their aggregation. For this purpose, we prepared two different types of liposomes/large unilamellar vesicles (LUVs), each carrying either a negative charge (DOPS) or neutral/zwitterionic (DOPC). Additionally, we performed cross-seeding experiment to observe any influence in the aggregation behavior of viral peptides. Amyloid aggregation of proteins advances via a nucleation-polymerization mechanism in which individual monomeric units assemble generating oligomers a.k.a. “critical seeds”(63, 64). These critical seeds have the ability to mature into beta-sheet rich fibrils as well as interact with monomers and promote their aggregation through a phenomenon referred to as self-seeding(65). Rationale behind Amylin protein selection is that it is a hallmark of type-2 diabetes (T2D), another well-studied protein misfolding disease(66). A positive correlation between diabetes and flaviviral infection has also been reported by many(67).

The aggregation kinetics was traced using ThT dye (same protocol described above) and the T-half (T_1/2_) was calculated which is the time at which half of the monomers has aggregated. We observed typical three prominent phases in kinetics curve of CA and 2K peptide characterized by an initial short “Lag phase” where the monomer nucleation occurs; followed by the exponential “Log phase” where oligomerization of monomers occurs giving rise to critical seeds as discussed above and final “plateau phase” where maturation of insoluble fibrils rich in beta-sheet takes place(68). Our results indicate a decrease in T-half (T_1/2_) of JEV CA when aggregated in the presence of LUVs (Figure 6 A,B). Interestingly, DOPS accelerated aggregation in CA more than DOPC. In the absence of LUVs, CA has a T_1/2_ of 6.43 h which was reduced to 6.14 h by DOPC and 4.86 h by DOPS (Figure 6B). This is probably because presence of biomembrane at an optimal concentration positively favors amyloid formation by condensing protein to increase effective concentration(24). They are also known to reduce the diffusional dimension and induce conformational changes in proteins, making them more susceptible to aggregation(24). On the other hand, cross-seeding with 2K and Amylin seeds at 10 μM concentration reduced T_1/2_ to 5.5 h and 4.62 h, respectively (Figure 6 C,D). This shows that 2K seeds and Amylin seeds were capable of interacting with CA monomers and enhance their rate of aggregation.

**Figure. 5:**
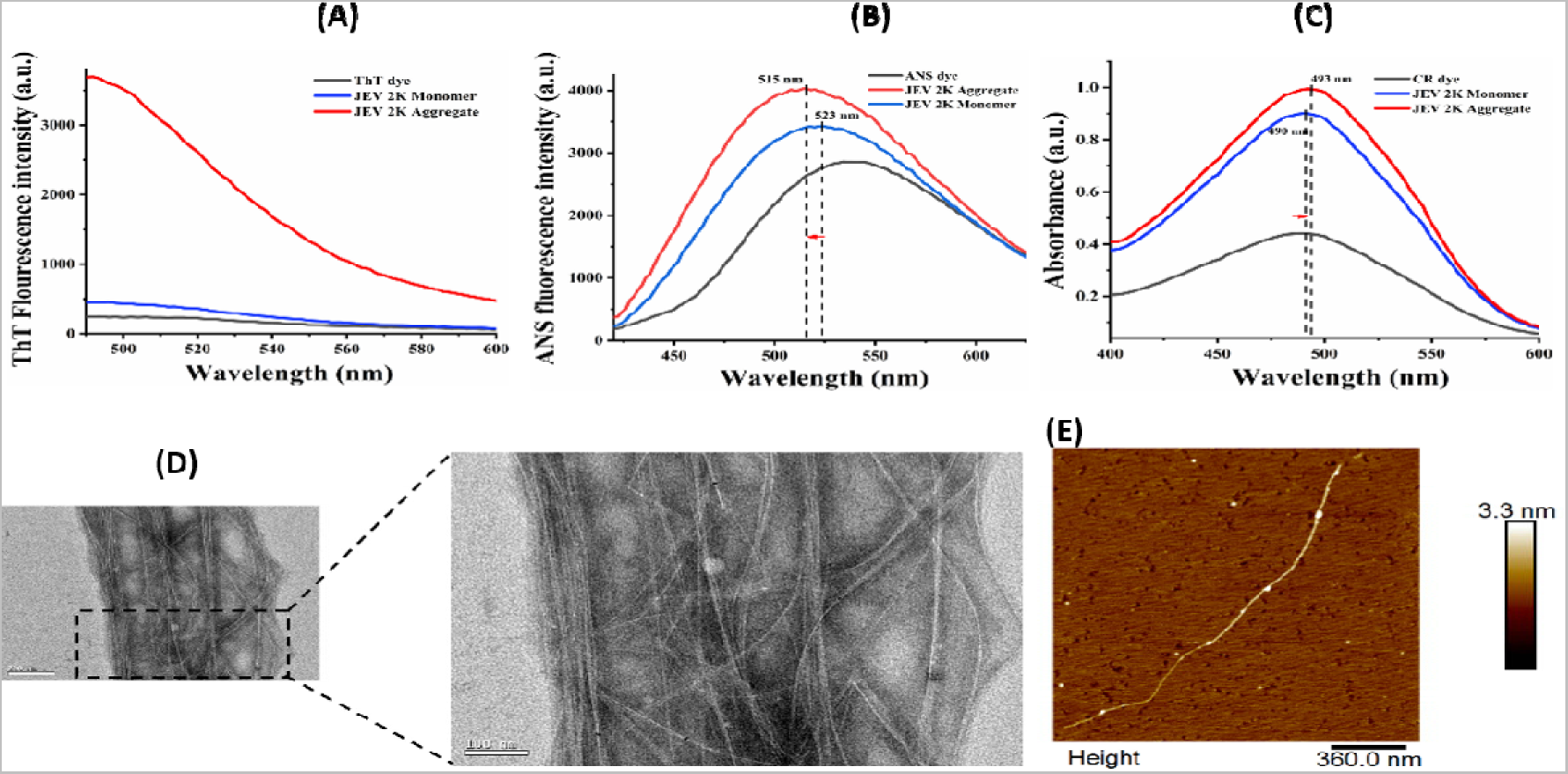
In vitro aggregation of the JEV 2K peptide. **(A)** Scans of ThT fluorescence spectra performed with 2K peptides at 1mg/ml concentration showing increased fluorescence intensity **(B)** ANS fluorescence emission spectra scan showing blue-shift **(C)** Absorption spectra scan of Congored dye bound capsid anchor peptides samples measured showing red-shift. TEM and AFM images of CA aggregates display typical amyloid fibril-like structures.

**Figure. 6:**
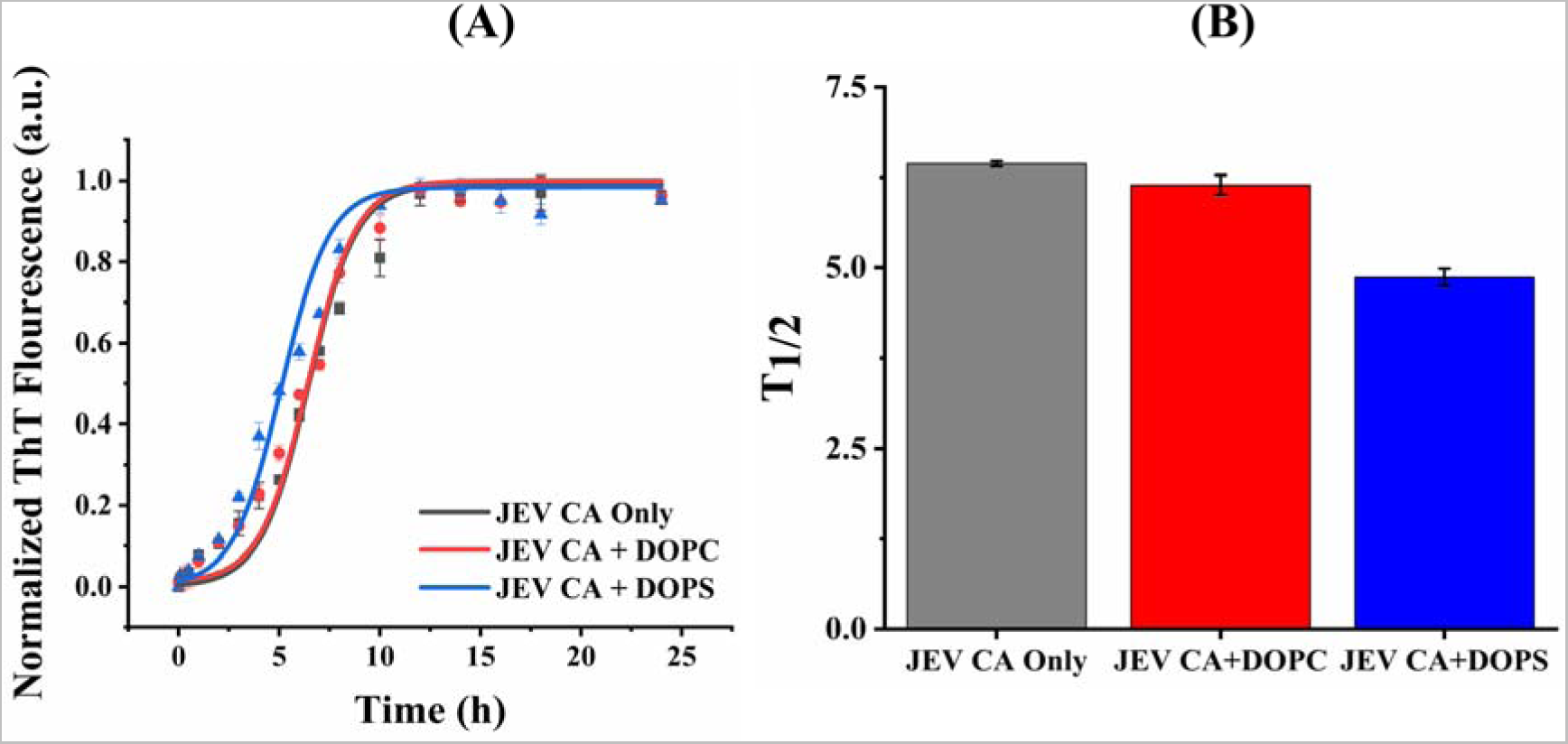
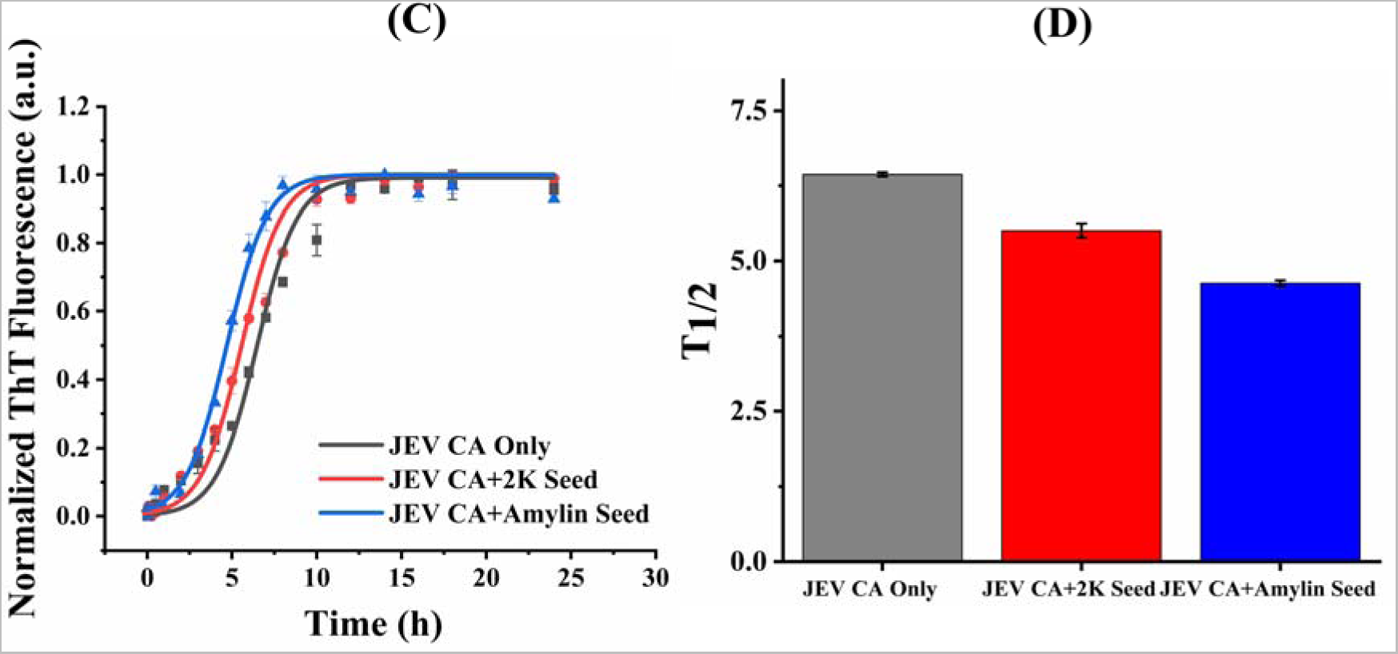
ThT aggregation kinetics. Plot and bar graph showing aggregation kinetics and T-half (T_1/2_) time of CA peptides in presence of **(A,B)** buffer only (Grey), zwitterionic DOPC (Red) and negatively charged DOPS (Blue) LUV. **(C,D)** buffer only (Grey), 2K seed (Red) and Amylin seed (Blue)

The JEV 2K aggregation kinetics has a T_1/2_ of 5.45 h (Figure 7A). This was reduced to 4.74 h by DOPC, 4.185 h by DOPS, 3.54 h by CA seed and 3.21 h by Amylin seeds, respectively (Figure 7 A-D). Overall results indicate that 2K has a relatively shorter kinetics compared to JEV CA in buffer, liposomes and critical-seeds. But the difference is not very significant and this could be attributed to the fact that both peptides are almost similar in length (JEV2K is longer by a single amino acid residue) and there is high similarity in their physicochemical properties and aggregate morphologies. Also, based on our observations, it is safe to say that JEV CA and 2K peptides can cross-seed and enhance aggregation of each other.

**Figure. 7:**
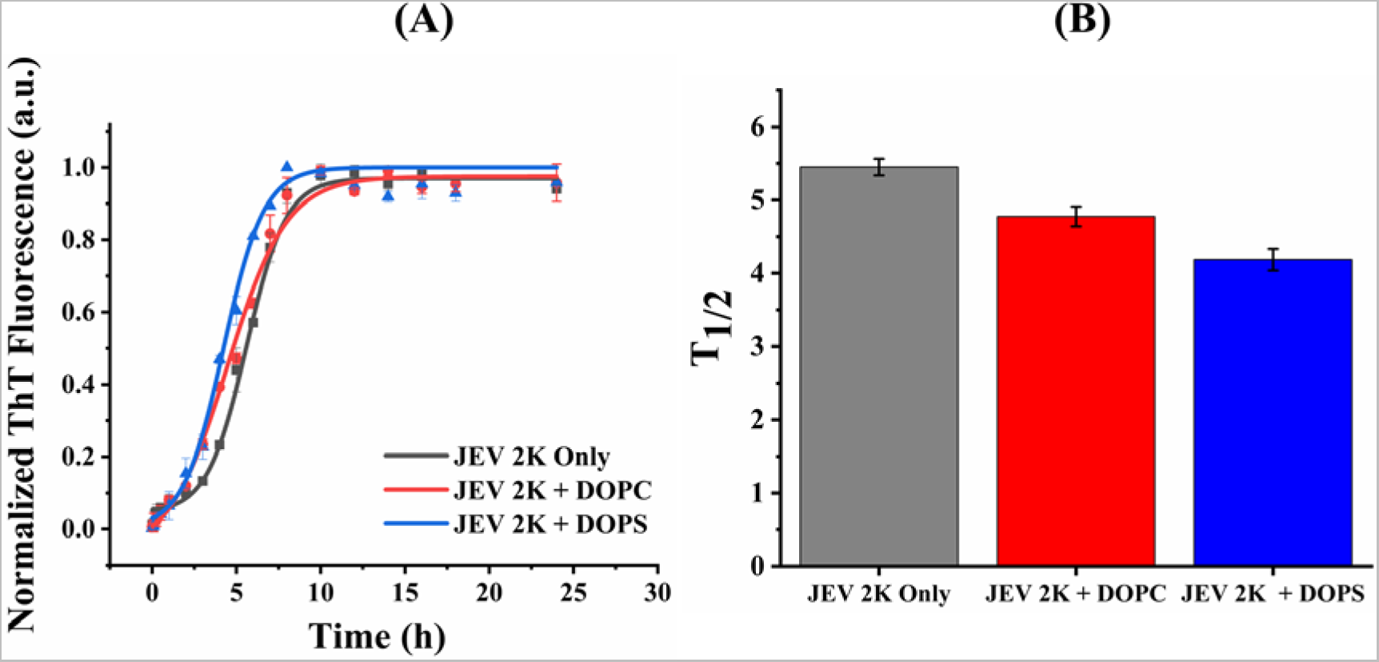
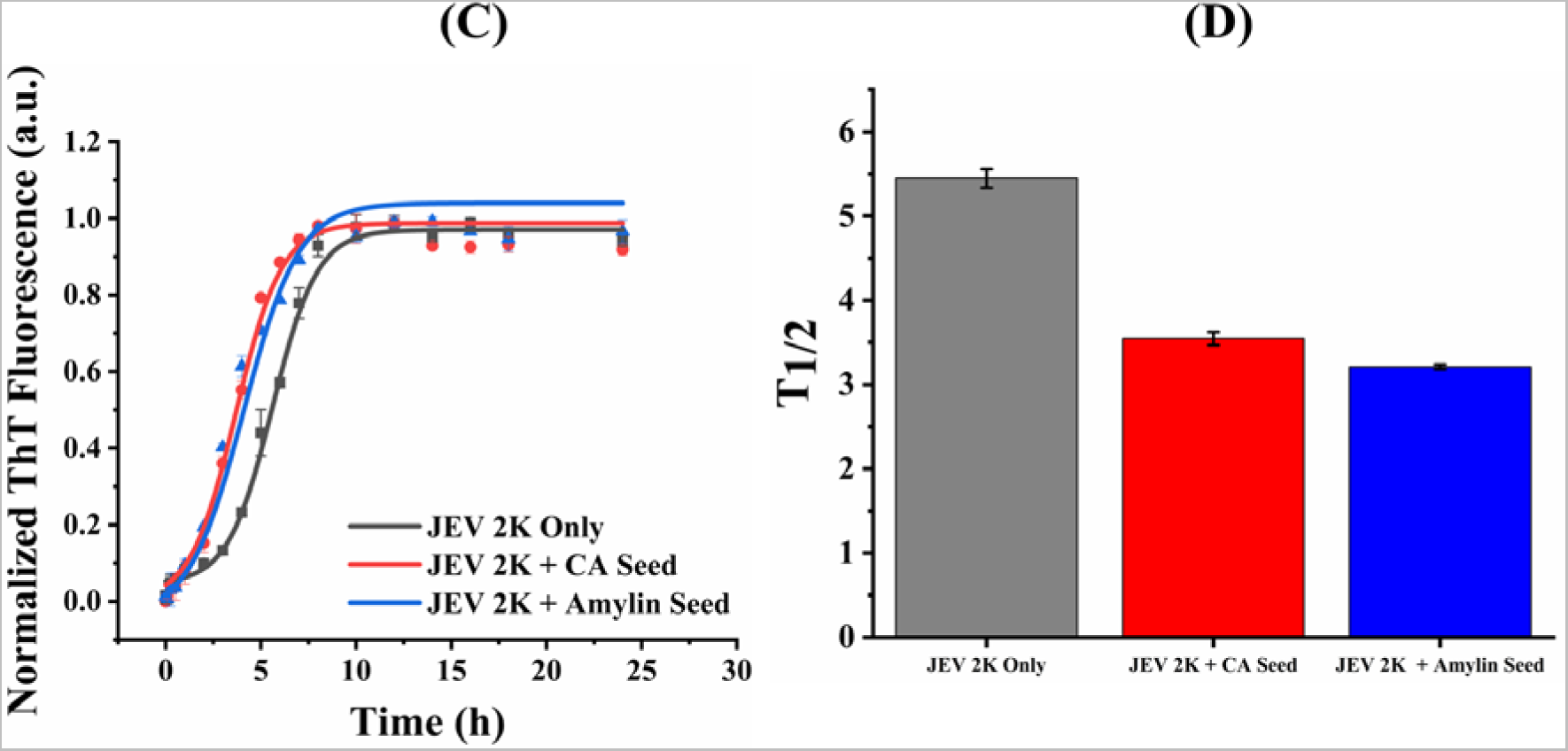
ThT aggregation kinetics. Plot and bar graph showing aggregation kinetics and T-half time (T_1/2_) of 2K peptides in presence of **(A,B)** buffer only (Grey), zwitterionic DOPC (Red) and negatively charged DOPS (Blue) LUV. **(C,D)** buffer only (Grey), 2K seed (Red) and Amylin seed (Blue)

### 3. JEV CA and 2K aggregates are cytotoxic to SH-SY5Y cells and damage human red blood cell (RBC) membranes

Pathogenic amyloid assemblies ranging from small oligomers to mature fibrillar structure has demonstrated the capacity to induce cytotoxicity(69–71). Recently, in our lab we have demonstrated that ZIKV CA and SARS-CoV proteins form cytotoxic amyloid fibrils too(20, 23). We were therefore curious to assess the possible cytotoxicity of the CA and 2K peptide aggregates in human derived neuroblastoma cells (SH-SY5Y) using 3-(4,5-dimethylthiazol-2-yl)-2,5-diphenyltetrazolium bromide (MTT) assay. The SH-SY5Y cells were incubated with five increasing concentrations of aggregates (5-500 µM) for 72 h duration. Monomers could not be used in this assay because of the poor solubility of synthetic peptides in buffer (20% DMSO used as per recommendation; Supplementary Figure 1-2). Figure 8 (A-C) shows results of the analysis of invitro results. The percentage cell viability reduced with increasing aggregate concentration of CA and 2K when compared to control (cells treated with 20mM sodium phosphate buffer, pH 7.4) suggesting their cytotoxic nature. From the lowest (5 µM) aggregate concentration to the highest (500µM) used in our reaction setup, the corresponding percent cell viability reduced from 99.47% to 74.59% for CA and 97.5% to 69.61% for 2K respectively (Figure 8 A,B). This indicates JEV 2K peptide forms slightly more cytotoxic amyloid fibrils than CA (Figure 8C).

**Figure 8.**
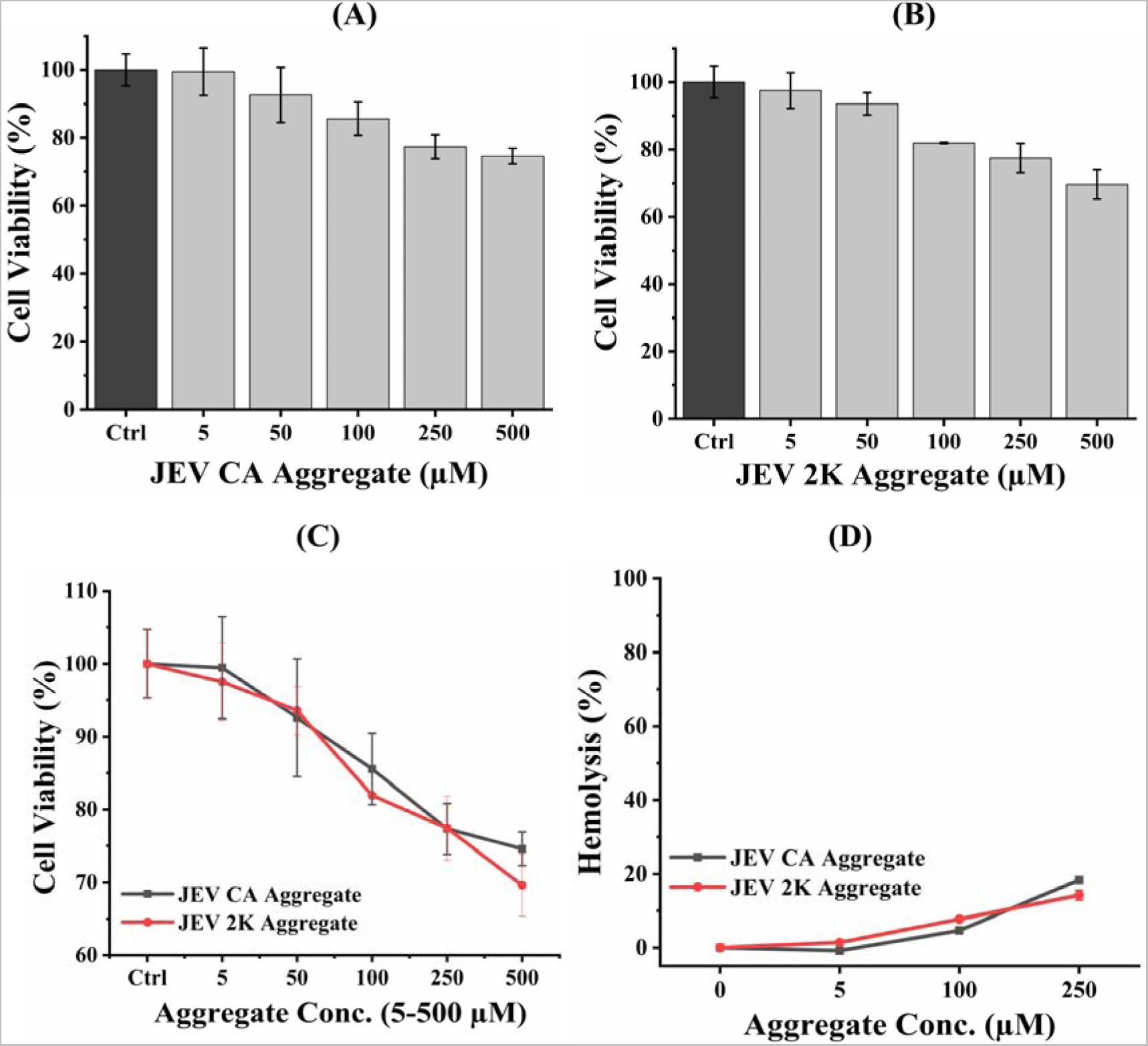
MTT assay and RBC haemolysis assay of JEV CA and 2K aggregates. **(A-C)** Bar graphs and plots showing percentage cell viability against various fibril concentrations of JEV CA (A), JEV 2K (B), combined [C; CA (Grey), 2K (Red)] aggregates (5-500 µM) relative to control. (D) Plot showing percentage haemolysis of human blood extracted RBCs against increasing concentration of CA (Black) and 2K (Red).

Based on the outcome of our cytotoxicity assay, we examined their effect on membrane lysis. We observed hemolysis results consistent with MTT assay where percent RBC lysis increased with increasing concentration of both CA and 2K peptides. The calculated percentage of hemolysis at highest aggregate concentration of 250µM was 18.36% and 14.18% for CA and 2K, respectively. Previous reports indicate that exposed hydrophobic regions in amyloid aggregates play a pivotal role in adhering to cell membranes and thereby exerting their detrimental impact on cells(72). The cytotoxic and RBC lytic effect of these two highly hydrophobic viral peptides could be attributed to this phenomenon.

## Conclusion

Protein misfolding into pathogenic aggregates are linked with several degenerative diseases. They typically arise from distinct amino acid sequences found within the fundamental framework of proteins and polypeptides. These sequences, referred to as aggregation-prone regions (APRs) possesses the ability to autonomously self-assemble into aggregates as well as engage with other protein via homologous interactions, leading to their conversion into beta-sheet rich insoluble structures. A significant number of these amyloids has been identified to play crucial roles in the progression of Alzheimer’s disease, Parkinsons disease and Type-2 diabetes. However, there exists a gap of our understanding between such pathological phenomenon and viral infections. JEV, a neurotropic flavivirus responsible for 20-30% case fatalities pose severe risk to over three billion people in endemic areas across the globe. Therefore, a comprehensive understanding of its infection mechanism is of utmost importance. In this study, where we analyzed JEV SA14 proteome for its aggregation propensities was inspired from series of recent reports in Zika virus, SARS CoV, Herpes Simplex virus-1, Epsten-Barr virus and Influenza-A virus (17–20, 23, 73). Our results indicated JEV to contain amyloid-prone regions across multiple functional and non-functional domains of its proteome. Two of its multifunctional transmembrane protein regions called “Capsid anchor” and “2K peptide” serve as linker between structural proteins (Capsid-2K-prM) and non-structural proteins (NS4A-2K-NS4B). In our in vitro analysis carried under physiological conditions, we observed that CA and 2K peptides are capable of aggregating and forming typical amyloid like structures. In the presence of simulated membranous environment and presence of critical-seeds, their aggregation rate enhanced greatly. Further, these aggregates could induce cell cytotoxicity probably through damaging membranes in contact. Thus, backed by our findings, we suggest that pathogenies of JE infection might be linked to aggregation of JEV proteins. Further in-depth analysis to gain insight into the molecular mechanism involved holds the potential to deepen our understanding of the virus as well as in designing of effective therapeutic strategies.

## Author Contributions

RG: Conception, design, and review of the manuscript. KUS: Acquisition and interpretation of data and writing of the manuscript. KUS, TB, DCT, DV: Acquisition of data.

## Acknowledgements

All the authors would like to thank IIT Mandi providing facilities. RG would like to acknowledge the Science and Engineering Research Board (SERB) (CRG/2019/005603), Department of Biotechnology, (BT/11/IYBA/2018/06), MHRD-SPARC (SPARC/2018-2019/P37/SL), and Indian Council of Medical Research (58/6/2020/PHA/BMS, and 52/04/2020/BIO/BMS). KUS is grateful to the ICMR for senior research fellowship Funding.

## References

1. Filgueira, L., and Lannes, N. (2019) Review of emerging japanese encephalitis virus: New aspects and concepts about entry into the brain and inter-cellular spreading. Pathogens. 10.3390/pathogens8030111

2. Schweitzer, B. K., Chapman, N. M., and Iwen, P. C. (2009) Overview of the Flaviviridae with an emphasis on the Japanese encephalitis group viruses. Lab. Med. 10.1309/LM5YWS85NJPCWESW

3. Kulkarni, R., Sapkal, G. N., Kaushal, H., and Mourya, D. T. (2018) Japanese Encephalitis: A Brief Review on Indian Perspectives. Open Virol. J. 10.2174/1874357901812010121

4. Ashraf, U., Ding, Z., Deng, S., Ye, J., Cao, S., and Chen, Z. (2021) Pathogenicity and virulence of Japanese encephalitis virus: Neuroinflammation and neuronal cell damage. Virulence. 10.1080/21505594.2021.1899674

5. Wang, X., Li, S. H., Zhu, L., Nian, Q. G., Yuan, S., Gao, Q., Hu, Z., Ye, Q., Li, X. F., Xie, D. Y., Shaw, N., Wang, J., Walter, T. S., Huiskonen, J. T., Fry, E. E., Qin, C. F., Stuart, D. I., and Rao, Z. (2017) Near-atomic structure of Japanese encephalitis virus reveals critical determinants of virulence and stability. Nat. Commun. 10.1038/s41467-017-00024-6

6. Fernandez-Garcia, M. D., Mazzon, M., Jacobs, M., and Amara, A. (2009) Pathogenesis of Flavivirus Infections: Using and Abusing the Host Cell. Cell Host Microbe. 10.1016/j.chom.2009.04.001

7. Dutta, K., Rangarajan, P. N., Vrati, S., and Basu, A. (2010) Japanese encephalitis: Pathogenesis, prophylactics and therapeutics. Curr. Sci.

8. Davis, E. H., Beck, A. S., Li, L., White, M. M., Greenberg, M. B., Thompson, J. K., Widen, S. G., Barrett, A. D. T., and Bourne, N. (2021) Japanese encephalitis virus live attenuated vaccine strains display altered immunogenicity, virulence and genetic diversity. npj Vaccines. 10.1038/s41541-021-00371-y

9. Aguzzi, A., and O’Connor, T. (2010) Protein aggregation diseases: Pathogenicity and therapeutic perspectives. Nat. Rev. Drug Discov. 10.1038/nrd3050

10. Hardy, J., and Selkoe, D. J. (2002) The amyloid hypothesis of Alzheimer’s disease: Progress and problems on the road to therapeutics. Science (80-.). 10.1126/science.1072994

11. Stefanis, L. (2012) α-Synuclein in Parkinson’s disease. Cold Spring Harb. Perspect. Med. 10.1101/cshperspect.a009399

12. Zheng, Z., and Diamond, M. I. (2012) Huntington disease and the huntingtin protein. in Progress in Molecular Biology and Translational Science, 10.1016/B978-0-12-385883-2.00010-2

13. Chen-Plotkin, A. S., Lee, V. M. Y., and Trojanowski, J. Q. (2010) TAR DNA-binding protein 43 in neurodegenerative disease. Nat. Rev. Neurol. 10.1038/nrneurol.2010.18

14. Head, M. W., and Ironside, J. W. (2012) Review: Creutzfeldt-Jakob disease: Prion protein type, disease phenotype and agent strain. Neuropathol. Appl. Neurobiol. 10.1111/j.1365-2990.2012.01265.x

15. Hayden, M. R., and Tyagi, S. C. (2001) “A” is for amylin and amyloid in type 2 diabetes mellitus. J. Pancreas

16. Taylor, J. P., Hardy, J., and Fischbeck, K. H. (2002) Toxic proteins in neurodegenerative disease. Science (80-.). 10.1126/science.1067122

17. Tiwari, D., Singh, V. K., Baral, B., Pathak, D. K., Jayabalan, J., Kumar, R., Tapryal, S., and Jha, H. C. (2021) Indication of Neurodegenerative Cascade Initiation by Amyloid-like Aggregate-Forming EBV Proteins and Peptide in Alzheimer’s Disease. ACS Chem. Neurosci. 10.1021/acschemneuro.1c00584

18. Chevalier, C., Al Bazzal, A., Vidic, J., Février, V., Bourdieu, C., Bouguyon, E., Le Goffic, R., Vautherot, J. F., Bernard, J., Moudjou, M., Noinville, S., Chich, J. F., Da Costa, B., Rezaei, H., and Delmas, B. (2010) PB1-F2 influenza A virus protein adopts a β-sheet conformation and forms amyloid fibers in membrane environments. J. Biol. Chem. 10.1074/jbc.M109.067710

19. Golovko, A. O., Koroleva, O. N., Tolstova, A. P., Kuz’mina, N. V., Dubrovin, E. V., and Drutsa, V. L. (2018) Aggregation of Influenza A Virus Nuclear Export Protein. Biochem. 10.1134/S0006297918110111

20. Saumya, K. U., Gadhave, K., Kumar, A., and Giri, R. (2021) Zika virus capsid anchor forms cytotoxic amyloid-like fibrils. Virology. 10.1016/j.virol.2021.04.010

21. Pham, C. L., Shanmugam, N., Strange, M., O’Carroll, A., Brown, J. W., Sierecki, E., Gambin, Y., Steain, M., and Sunde, M. (2019) Viral M45 and necroptosis[associated proteins form heteromeric amyloid assemblies. EMBO Rep. 10.15252/embr.201846518

22. Zlotnick, A., Aldrich, R., Johnson, J. M., Ceres, P., and Young, M. J. (2000) Mechanism of capsid assembly for an icosahedral plant virus. Virology. 10.1006/viro.2000.0619

23. Bhardwaj, T., Gadhave, K., Kapuganti, S. K., Kumar, P., Faidon Brotzakis, Z., Udit Saumya, K., Nayak, N., Kumar, A., Garg, N., Vendruscolo, M., and Giri, R. (2021) Amyloidogenic proteins in the SARS-CoV and SARS-CoV-2 proteomes. bioRxiv

24. Terakawa, M. S., Lin, Y., Kinoshita, M., Kanemura, S., Itoh, D., Sugiki, T., Okumura, M., Ramamoorthy, A., and Lee, Y. H. (2018) Impact of membrane curvature on amyloid aggregation. Biochim. Biophys. Acta - Biomembr. 10.1016/j.bbamem.2018.04.012

25. Emily, M., Talvas, A., and Delamarche, C. (2013) MetAmyl: A METa-predictor for AMYLoid proteins. PLoS One. 10.1371/journal.pone.0079722

26. Garbuzynskiy, S. O., Lobanov, M. Y., and Galzitskaya, O. V. (2009) FoldAmyloid: A method of prediction of amyloidogenic regions from protein sequence. Bioinformatics. 26, 326–332

27. Fernandez-Escamilla, A. M., Rousseau, F., Schymkowitz, J., and Serrano, L. (2004) Prediction of sequence-dependent and mutational effects on the aggregation of peptides and proteins. Nat. Biotechnol. 22, 1302–1306

28. Conchillo-Solé, O., de Groot, N. S., Avilés, F. X., Vendrell, J., Daura, X., and Ventura, S. (2007) AGGRESCAN: A server for the prediction and evaluation of “hot spots” of aggregation in polypeptides. BMC Bioinformatics. 10.1186/1471-2105-8-65

29. Gasior, P., and Kotulska, M. (2014) FISH Amyloid - a new method for finding amyloidogenic segments in proteins based on site specific co-occurence of aminoacids. BMC Bioinformatics. 10.1186/1471-2105-15-54

30. Gadhave, K., Bhardwaj, T., Uversky, V. N., Vendruscolo, M., and Giri, R. (2021) The signal peptide of the amyloid precursor protein forms amyloid-like aggregates and enhances Aβ42 aggregation. Cell Reports Phys. Sci. 10.1016/j.xcrp.2021.100599

31. Saumya, K. U., Kumar, D., Kumar, P., and Giri, R. (2020) Unlike dengue virus, the conserved 14–23 residues in N-terminal region of Zika virus capsid is not involved in lipid interactions. Biochim. Biophys. Acta - Biomembr. 10.1016/j.bbamem.2020.183440

32. Weids, A. J., Ibstedt, S., Tamás, M. J., and Grant, C. M. (2016) Distinct stress conditions result in aggregation of proteins with similar properties. Sci. Rep. 10.1038/srep24554

33. Rousseau, F., Schymkowitz, J., and Serrano, L. (2006) Protein aggregation and amyloidosis: Confusion of the kinds? Curr. Opin. Struct. Biol. 10.1016/j.sbi.2006.01.011

34. Mukhopadhyay, S., Kuhn, R. J., and Rossmann, M. G. (2005) A structural perspective of the Flavivirus life cycle. Nat. Rev. Microbiol. 10.1038/nrmicro1067

35. Oliveira, E. R. A., Mohana-Borges, R., de Alencastro, R. B., and Horta, B. A. C. (2017) The flavivirus capsid protein: Structure, function and perspectives towards drug design. Virus Res. 227, 115–123

36. Poonsiri, T., Wright, G. S. A., Solomon, T., and Antonyuk, S. V. (2019) Crystal structure of the Japanese encephalitis virus capsid protein. Viruses. 10.3390/v11070623

37. Luca, V. C., AbiMansour, J., Nelson, C. A., and Fremont, D. H. (2012) Crystal Structure of the Japanese Encephalitis Virus Envelope Protein. J. Virol. 10.1128/jvi.06072-11

38. Hsieh, S. C., Wu, Y. C., Zou, G., Nerurkar, V. R., Shi, P. Y., and Wang, W. K. (2014) Highly conserved residues in the helical domain of dengue virus type 1 precursor membrane protein are involved in assembly, precursor membrane (prM) protein cleavage, and entry. J. Biol. Chem. 10.1074/jbc.M114.610428

39. Rastogi, M., Sharma, N., and Singh, S. K. (2016) Flavivirus NS1: A multifaceted enigmatic viral protein. Virol. J. 10.1186/s12985-016-0590-7

40. Zhou, D., Jia, F., Li, Q., Zhang, L., Chen, Z., Zhao, Z., Cui, M., Song, Y., Chen, H., Cao, S., and Ye, J. (2018) Japanese Encephalitis Virus NS1′ Protein Antagonizes Interferon Beta Production. Virol. Sin. 10.1007/s12250-018-0067-5

41. Takamatsu, Y., Morita, K., and Hayasaka, D. (2015) A Single Amino Acid Substitution in the NS2A Protein of Japanese Encephalitis Virus Affects Virus Propagation In Vitro but Not In Vivo. J. Virol. 10.1128/jvi.00370-15

42. Jan, L. R., Yang, C. S., Trent, D. W., Falgout, B., and Lai, C. J. (1995) Processing of Japanese encephalitis virus non-structural proteins: NS2B-NS3 complex and heterologous proteases. J. Gen. Virol. 10.1099/0022-1317-76-3-573

43. León-Juárez, M., Martínez-Castillo, M., Shrivastava, G., García-Cordero, J., Villegas-Sepulveda, N., Mondragón-Castelán, M., Mondragón-Flores, R., and Cedillo-Barrón, L. (2016) Recombinant Dengue virus protein NS2B alters membrane permeability in different membrane models. Virol. J. 10.1186/s12985-015-0456-4

44. Yamashita, T., Unno, H., Mori, Y., Tani, H., Moriishi, K., Takamizawa, A., Agoh, M., Tsukihara, T., and Matsuura, Y. (2008) Crystal structure of the catalytic domain of Japanese encephalitis virus NS3 helicase/nucleoside triphosphatase at a resolution of 1.8 Å. Virology. 10.1016/j.virol.2007.12.018

45. Bhardwaj, T., Saumya, K. U., Kumar, P., Sharma, N., Gadhave, K., Uversky, V. N., and Giri, R. (2020) Japanese encephalitis virus – exploring the dark proteome and disorder–function paradigm. FEBS J. 10.1111/febs.15427

46. Shiryaev, S. A., Chernov, A. V., Aleshin, A. E., Shiryaeva, T. N., and Strongin, A. Y. (2009) NS4A regulates the ATPase activity of the NS3 helicase: A novel cofactor role of the non-structural protein NS4A from West Nile virus. J. Gen. Virol. 10.1099/vir.0.012864-0

47. Li, X. D., Ye, H. Q., Deng, C. L., Liu, S. Q., Zhang, H. L., Shang, B. Di, Shi, P. Y., Yuan, Z. M., and Zhang, B. (2015) Genetic interaction between NS4A and NS4B for replication of Japanese encephalitis virus. J. Gen. Virol. 10.1099/vir.0.000044

48. Lin, C., Amberg, S. M., Chambers, T. J., and Rice, C. M. (1993) Cleavage at a novel site in the NS4A region by the yellow fever virus NS2B-3 proteinase is a prerequisite for processing at the downstream 4A/4B signalase site. J. Virol.

49. Lu, G., and Gong, P. (2013) Crystal Structure of the Full-Length Japanese Encephalitis Virus NS5 Reveals a Conserved Methyltransferase-Polymerase Interface. PLoS Pathog. 10.1371/journal.ppat.1003549

50. Lin, R.-J., Chang, B.-L., Yu, H.-P., Liao, C.-L., and Lin, Y.-L. (2006) Blocking of Interferon-Induced Jak-Stat Signaling by Japanese Encephalitis Virus NS5 through a Protein Tyrosine Phosphatase-Mediated Mechanism. J. Virol. 10.1128/jvi.02714-05

51. Rana, J., Slon Campos, J. L., Leccese, G., Francolini, M., Bestagno, M., Poggianella, M., and Burrone, O. R. (2018) Role of Capsid Anchor in the Morphogenesis of Zika Virus. J. Virol. 10.1128/jvi.01174-18

52. Burrone, O. R., Campos, J. L. S., Poggianella, M., and Rana, J. (2020) Impact of Capsid Anchor Length and Sequential Processing on the Assembly and Infectivity of Dengue Virus. Proceedings. 50, 32

53. Blazevic, J., Rouha, H., Bradt, V., Heinz, F. X., and Stiasny, K. (2016) Membrane Anchors of the Structural Flavivirus Proteins and Their Role in Virus Assembly. J. Virol. 90, 6365–6378

54. Biancalana, M., and Koide, S. (2010) Molecular mechanism of Thioflavin-T binding to amyloid fibrils. Biochim. Biophys. Acta - Proteins Proteomics. 10.1016/j.bbapap.2010.04.001

55. Hawe, A., Sutter, M., and Jiskoot, W. (2008) Extrinsic fluorescent dyes as tools for protein characterization. Pharm. Res. 10.1007/s11095-007-9516-9

56. Frid, P., Anisimov, S. V., and Popovic, N. (2007) Congo red and protein aggregation in neurodegenerative diseases. Brain Res. Rev. 10.1016/j.brainresrev.2006.08.001

57. Gadhave, K., and Giri, R. (2020) Amyloid formation by intrinsically disordered trans-activation domain of cMyb. Biochem. Biophys. Res. Commun. 10.1016/j.bbrc.2020.01.110

58. Kumar, S., Verma, A., Yadav, P., Dubey, S. K., Azhar, E. I., Maitra, S. S., and Dwivedi, V. D. (2022) Molecular pathogenesis of Japanese encephalitis and possible therapeutic strategies. Arch. Virol. 10.1007/s00705-022-05481-z

59. Lobigs, M., Lee, E., Ng, M. L., Pavy, M., and Lobigs, P. (2010) A flavivirus signal peptide balances the catalytic activity of two proteases and thereby facilitates virus morphogenesis. Virology. 401, 80–89

60. Barnard, T. R., Abram, Q. H., Lin, Q. F., Wang, A. B., and Sagan, S. M. (2021) Molecular Determinants of Flavivirus Virion Assembly. Trends Biochem. Sci. 10.1016/j.tibs.2020.12.007

61. Zou, G., Puig-Basagoiti, F., Zhang, B., Qing, M., Chen, L., Pankiewicz, K. W., Felczak, K., Yuan, Z., and Shi, P. Y. (2009) A single-amino acid substitution in West Nile virus 2K peptide between NS4A and NS4B confers resistance to lycorine, a flavivirus inhibitor. Virology. 10.1016/j.virol.2008.11.003

62. Aisenbrey, C., Borowik, T., Byström, R., Bokvist, M., Lindström, F., Misiak, H., Sani, M. A., and Gröbner, G. (2008) How is protein aggregation in amyloidogenic diseases modulated by biological membranes? in European Biophysics Journal, 10.1007/s00249-007-0237-0

63. Morales, R., Estrada, L. D., Diaz-Espinoza, R., Morales-Scheihing, D., Jara, M. C., Castilla, J., and Soto, C. (2010) Molecular cross talk between misfolded proteins in animal models of alzheimer’s and prion diseases. J. Neurosci. 10.1523/JNEUROSCI.5924-09.2010

64. Ren, B., Zhang, Y., Zhang, M., Liu, Y., Zhang, D., Gong, X., Feng, Z., Tang, J., Chang, Y., and Zheng, J. (2019) Fundamentals and Introductory of Cross-seeding of Amyloid Proteins. J. Mater. Chem. B

65. Soto, C., and Pritzkow, S. (2018) Protein misfolding, aggregation, and conformational strains in neurodegenerative diseases. Nat. Neurosci. 10.1038/s41593-018-0235-9

66. Hassan, S., White, K., and Terry, C. (2022) Linking hIAPP misfolding and aggregation with type 2 diabetes mellitus: a structural perspective. Biosci. Rep. 10.1042/BSR20211297

67. Sorenson, A., Owens, L., Caltabiano, M., Cadet-James, Y., Hall, R., Govan, B., and Clancy, P. (2016) The impact of prior flavivirus infections on the development of type 2 diabetes among the Indigenous Australians. Am. J. Trop. Med. Hyg. 10.4269/ajtmh.15-0727

68. Lee, C. C., Nayak, A., Sethuraman, A., Belfort, G., and McRae, G. J. (2007) A three-stage kinetic model of amyloid fibrillation. Biophys. J. 10.1529/biophysj.106.098608

69. Marshall, K. E., Marchante, R., Xue, W. F., and Serpell, L. C. (2014) The relationship between amyloid structure and cytotoxicity. Prion. 10.4161/pri.28860

70. Yoshiike, Y., Akagi, T., and Takashima, A. (2007) Surface structure of amyloid-β fibrils contributes to cytotoxicity. Biochemistry. 10.1021/bi700455c

71. Bystrenova, E., Bednarikova, Z., Barbalinardo, M., Albonetti, C., Valle, F., and Gazova, Z. (2019) Amyloid fragments and their toxicity on neural cells. Regen. Biomater. 10.1093/rb/rbz007

72. Andreasen, M., Lorenzen, N., and Otzen, D. (2015) Interactions between misfolded protein oligomers and membranes: A central topic in neurodegenerative diseases? Biochim. Biophys. Acta - Biomembr. 10.1016/j.bbamem.2015.01.018

73. Singh, V. K., Kumar, S., and Tapryal, S. (2020) Aggregation Propensities of Herpes Simplex Virus-1 Proteins and Derived Peptides: An in Silico and in Vitro Analysis. ACS Omega. 10.1021/acsomega.0c00730

